# Demography and mating system shape the genome-wide impact of purifying selection in *Arabis alpina*

**DOI:** 10.1101/127209

**Authors:** Benjamin Laenen, Andrew Tedder, Michael D. Nowak, Per Toräng, Jörg Wunder, Stefan Wötzel, Kim A. Steige, Yiannis Kourmpetis, Thomas Odong, Andreas D. Drouzas, Marco Bink, Jon Ågren, George Coupland, Tanja Slotte

**Author notes:** B.L. and A.T. contributed equally to this work. Authors for correspondence, email: Jon Ågren; George Coupland, Tanja Slotte.

## Abstract

Plant mating systems have profound effects on levels and structuring of genetic variation, and can affect the impact of natural selection. While theory predicts that intermediate outcrossing rates may allow plants to prevent accumulation of deleterious alleles, few studies have empirically tested this prediction using genomic data. Here, we study the effect of mating system on purifying selection by conducting population genomic analyses on whole-genome resequencing data from 38 European individuals of the arctic-alpine crucifer *Arabis alpina*. We find that outcrossing and mixed-mating populations maintain genetic diversity at similar levels, whereas highly self-fertilizing Scandinavian *A. alpina* show a strong reduction in genetic diversity, most likely as a result of a postglacial colonization bottleneck. We further find evidence for accumulation of genetic load in highly self-fertilizing populations, whereas the genome-wide impact of purifying selection does not differ greatly between mixed-mating and outcrossing populations. Our results demonstrate that intermediate levels of outcrossing may allow efficient selection against harmful alleles whereas demographic effects can be important for relaxed purifying selection in highly selfing populations. Thus, both mating system and demography shape the impact of purifying selection on genomic variation in *A. alpina*. These results are important for an improved understanding of the evolutionary consequences of mating system variation and the maintenance of mixed-mating strategies.

**Significance:** Intermediate outcrossing rates are theoretically predicted to maintain effective selection against harmful alleles, but few studies have empirically tested this prediction using genomic data. We used whole-genome resequencing data from alpine rock-cress to study how genetic variation and purifying selection vary with mating system. We find that populations with intermediate outcrossing rates have similar levels of genetic diversity as outcrossing populations, and that purifying selection against harmful alleles is efficient in mixed-mating populations. In contrast, self-fertilizing populations from Scandinavia have strongly reduced genetic diversity, and accumulate harmful mutations, likely as a result of demographic effects of postglacial colonization. Our results suggest that mixed-mating populations can avoid the negative evolutionary consequences of high self-fertilization rates.

## Introduction

Flowering plants show a great deal of variation in their reproductive modes, and variation in outcrossing rate is particularly common. While approximately 50% of flowering plants are predominantly outcrossing, a substantial proportion (35-40%) undergo intermediate levels of outcrossing, whereas only 10-15% are predominantly self-fertilizing (selfing) (1, 2). Whether mixed-mating is evolutionarily stable or represents a transitional stage has long been debated (1, 3). Classic population genetic models predict that only high selfing and high outcrossing rates are evolutionarily stable strategies (4). However, mixed-mating can be stable in ecologically more realistic models, such as those that account for reduced outcross pollen success with increased selfing (1). Moreover, population genetic models that incorporate linkage indicate that mixed-mating populations may avoid the reduced efficacy of selection associated with high selfing rates (5-7).

While selfing can be favored due to its genetic transmission advantage, and because it can confer reproductive assurance, it also has marked population genetic consequences that might contribute to the long-term demise of highly selfing lineages (6). For instance, highly selfing populations are expected to have a reduced effective population size (*N*_*e*_), due to the direct effect of inbreeding (8, 9). Demographic processes such as frequent extinction and recolonization of local subpopulations (10) or founder events associated with the shift to selfing (6) can reduce genetic variation in selfers to an even greater extent. Self-fertilization is also expected to lead to elevated linkage disequilibrium, which means that background selection and other forms of linked selection can reduce genetic variation genome-wide, further reducing *N*_*e*_ in selfers (7, 11, 12).

Because the strength of selection scales with *N*_*e*_ (13), natural selection is expected to be less efficient in selfers than in outcrossers. Selfers are therefore expected to accumulate weakly deleterious mutations at a higher rate than outcrossers (14). However, selfing also increases homozygosity, exposing recessive mutations to selection. This is expected to result in purging of recessive deleterious mutations, unless selfing is associated with a strong reduction in *N*_*e*_ which renders such purging ineffective (15-17).

While there is accumulating evidence for relaxed selection on weakly deleterious mutations in highly selfing lineages (5, 18-22), several fundamental questions on the evolutionary genomic consequences of mating systems remain unanswered. First, it is unclear whether selfing generally results in purging of recessive deleterious alleles or whether this effect is possibly overridden by demographic effects. Second, it remains unclear whether intermediate levels of outcrossing allow plants to avoid the negative evolutionary genetic consequences of high selfing rates. Specifically, while theory predicts that intermediate levels of self-fertilization could be sufficient to achieve purging of deleterious alleles (16) and prevent accumulation of mildly deleterious alleles (23), few empirical genome-wide studies have examined the selective consequences of mixed-mating in plants (but see (24)).

The broadly distributed, arctic-alpine perennial herb *Arabis alpina* (Brassicaceae) is a promising plant system in which to address the impact of variation in outcrossing rates on genome-wide genetic variation and efficiency of selection. This species harbors populations that express a range of mating strategies from self-incompatible outcrossing (25) through mixed-mating to autonomous selfing (25-27). The colonization history of *A. alpina* has already begun to be characterized (26, 28-30), which facilitates interpretation of global patterns of polymorphism. Finally, the availability of a genome assembly of *A. alpina* (31) greatly facilitates population genomic studies.

In this study, we investigate the effects of mating system and demography on the efficacy of selection in *A. alpina* by population genomic analyses of whole-genome resequencing data from outcrossing, mixed-mating and highly selfing populations. We first investigate population structure and test whether populations with higher selfing rates have lower levels of genetic diversity, and then test whether higher selfing rates are associated with relaxed selection against weakly deleterious mutations and purging of strongly deleterious mutations. To do this, we use genome-wide allele frequency distributions at nonsynonymous and synonymous sites to estimate the fraction of weakly deleterious and strongly harmful new nonsynonymous mutations. We further test whether higher rates of selfing are associated with an increase in the frequency of derived alleles with major effects on gene integrity, which would suggest relaxed purifying selection. Finally, we compare two genomic proxies of genetic load, the reduction in mean fitness of a population caused by deleterious variation (32,33), among outcrossing, mixed-mating and highly self-fertilizing populations. Our results are important for an improved understanding of the population genetic consequences of mating system variation.

## Results

### Sequencing and single nucleotide polymorphism

We sampled 38 *A. alpina* individuals, with targeted sampling of 2-5 individuals from 12 geographical sites harboring self-incompatible outcrossing populations (Greece and Italy), populations with intermediate outcrossing rates (France and Spain), and highly selfing populations (Scandinavia), in addition to a sample of single individuals from 5 additional geographic locations across the European range of *A. alpina* (SI Appendix, Table S1). Progeny-array based outcrossing estimates have shown that populations from Scandinavia are highly selfing (up to ∼10% outcrossing) whereas intermediate outcrossing rates have been estimated for two French and Spanish populations (∼20% and ∼ 18%, respectively) (SI Appendix, SI Text). We further verified that Greek and Italian individuals produced no offspring after forced selfing in the greenhouse. The distribution of genomic runs of homozygosity (ROHs) further supported mating system variation among the populations studied here (SI Appendix, SI Text).

Each individual was resequenced to high coverage (average 26X, range 16-45X) using Illumina short-read technology. We called single nucleotide polymorphisms (SNPs) and applied stringent filtering criteria to identify a total of 1,514,615 high-quality SNPs, of which 98,564 were 0-fold degenerate nonsynonymous (i.e. sites at which any mutation will result in a nonsynonymous change) with a mean nucleotide diversity (π) of 0.0027 and 65,821 were 4-fold degenerate synonymous (i.e. synonymous sites at which any mutation will result in a synonymous change) with a mean nucleotide diversity of 0.0102 (SI Appendix, Table S2).

### Population structure has a strong geographic component

We analyzed population structure with two model-based Bayesian clustering approaches, implemented in the software fastSTRUCTURE (34) and TESS3 (35), based on 25,505 4-fold synonymous SNPs (see Methods). Both methods, as well as principal component analysis gave very similar results, and supported the presence of five clusters (Fig. 1; SI Appendix, Fig. S1), in good agreement with previous analyses of population structure in *A. alpina* (29). These clusters correspond to a central European population of mixed-mating individuals from France, Germany, Poland and Switzerland, a northern European population of highly self-fertilizing individuals from Sweden, Norway and Iceland, a mixed-mating population containing individuals from Spain and Madeira, and two outcrossing populations representing individuals from Italy and Greece, respectively (Fig. 1). Subsequent population genetic analyses are presented separately for regional population samples from each of these geographical regions.

**Figure 1.**
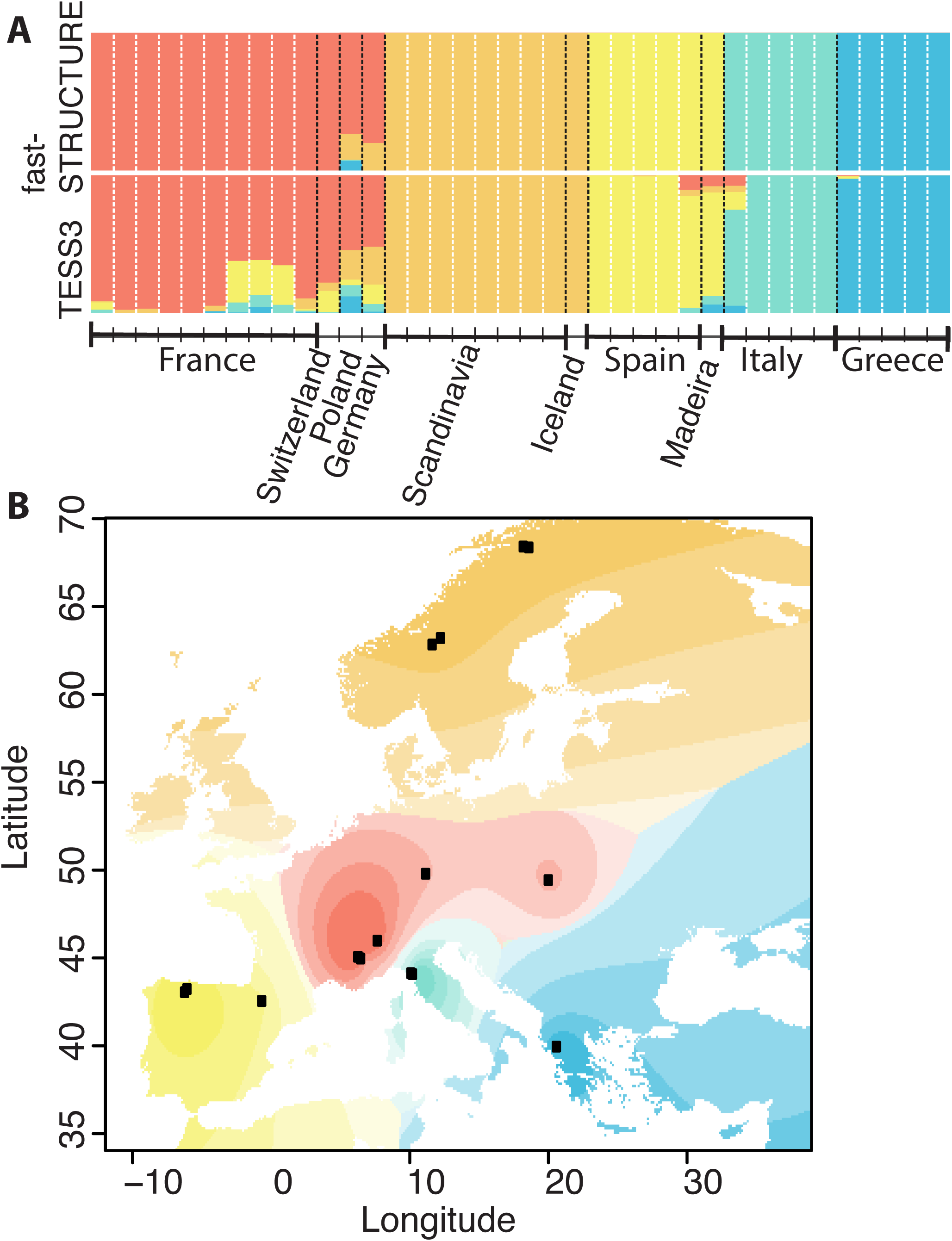
Bayesian clustering analysis supports a strong geographic component to population structure in European *A. alpina*. A. Ancestry proportions for K=5 correspond closely to geographical sampling locations. B. Geographic interpolation of genetic structure across Europe based on TESS3 results for K=5.

### Genetic diversity is maintained in mixed-mating but not in highly selfing populations

For each regional population, we quantified nucleotide diversity at three categories of sites: 4-fold degenerate sites, 0-fold degenerate sites and intergenic sites in regions of low gene density and high recombination rate (Table 1, SI Appendix, Table S3). At all three categories of sites, levels of nucleotide diversity (π) varied by an order of magnitude among regional populations (Table 1, SI Appendix, Table S3). The outcrossing Greek population was the most genetically diverse (4-fold synonymous diversity π_S_ = 0.008), whereas the highly self-fertilizing Scandinavian populations had very low nucleotide diversity (π_S_ =0.0002; Table 1). Levels of nucleotide diversity were intermediate and of a similar magnitude in both outcrossing Italian and mixed-mating French and Spanish populations (Table 1), suggesting that in *A. alpina,* mixed-mating populations maintain similar levels of genetic diversity as outcrossing populations. Similar patterns were seen for 0-fold nonsynonymous sites, although the relative reduction in diversity in Scandinavia was less severe for 0-fold sites than for 4-fold sites (SI Appendix, Table S3). This resulted in a markedly elevated ratio of nonsynonymous to synonymous nucleotide diversity (*π*_*N*_/*π*_*S*_) in Scandinavia (Table 1).

**Table 1.** Population genetic summary statistics for regional populations.

| Population<sup>1</sup> | S<sub>s</sub><sup>2</sup> | π<sub>s</sub><sup>3</sup> | π<sub>N</sub>/π<sub>s</sub><sup>4</sup> |
| --- | --- | --- | --- |
| Greece (5) | 30920 | 0.0080 | 0.2740 |
| Italy (5) | 17971 | 0.0054 | 0.2820 |
| Spain (5) | 15852 | 0.0046 | 0.2911 |
| France (10) | 19661 | 0.0052 | 0.2764 |
| Scandinavia (8) | 589 | 0.0002 | 0.3936 |

<sup>1</sup>Regional population (*n*)
<sup>2</sup>Segregating four-fold synonymous SNPs.
<sup>3</sup>Mean four-fold synonymous nucleotide diversity.
<sup>4</sup>Mean 0-fold degenerate nonsynonymous/4-fold degenerate synonymous polymorphism.

### Selfing, but not mixed-mating, is associated with relaxed purifying selection

To test whether the impact of purifying selection varied with mating system, we estimated the distribution of negative fitness effects (DFE) of new nonsynonymous mutations using DFE-alpha (37) (See Methods for details). This method can detect changes in selection in association with plant mating system shifts, including purging of strongly deleterious mutations (38), and corrects for effects of demographic changes on allele frequency spectra using a simple population size change model (37). We summarized the strength of selection, defined as the product of the effective population size *N*_*e*_ and the selection coefficient *s*, in three bins, ranging from nearly neutral to strongly deleterious (0<*N*_*e*_*s*<1; 1<*N*_*e*_*s*<10; *N*_*e*_*s*>10). While the DFE of the mixed-mating Spanish and outcrossing Greek populations differed, there were no significant differences in the DFE of the outcrossing Italian population and the mixed-mating Spanish and French populations (Fig. 2A). We detected no signature of purging of strongly deleterious mutations in mixed-mating populations based on DFE analyses (Fig. 2A), as the inferred proportion of strongly deleterious new nonsynonymous mutations was not generally higher in the mixed-mating French and Spanish than in the outcrossing Greek and Italian populations. This suggests that *A. alpina* populations undergoing as much as 80% selfing do not show evidence of relaxed purifying selection or increased purging of recessive deleterious mutations.

**Figure 2.**
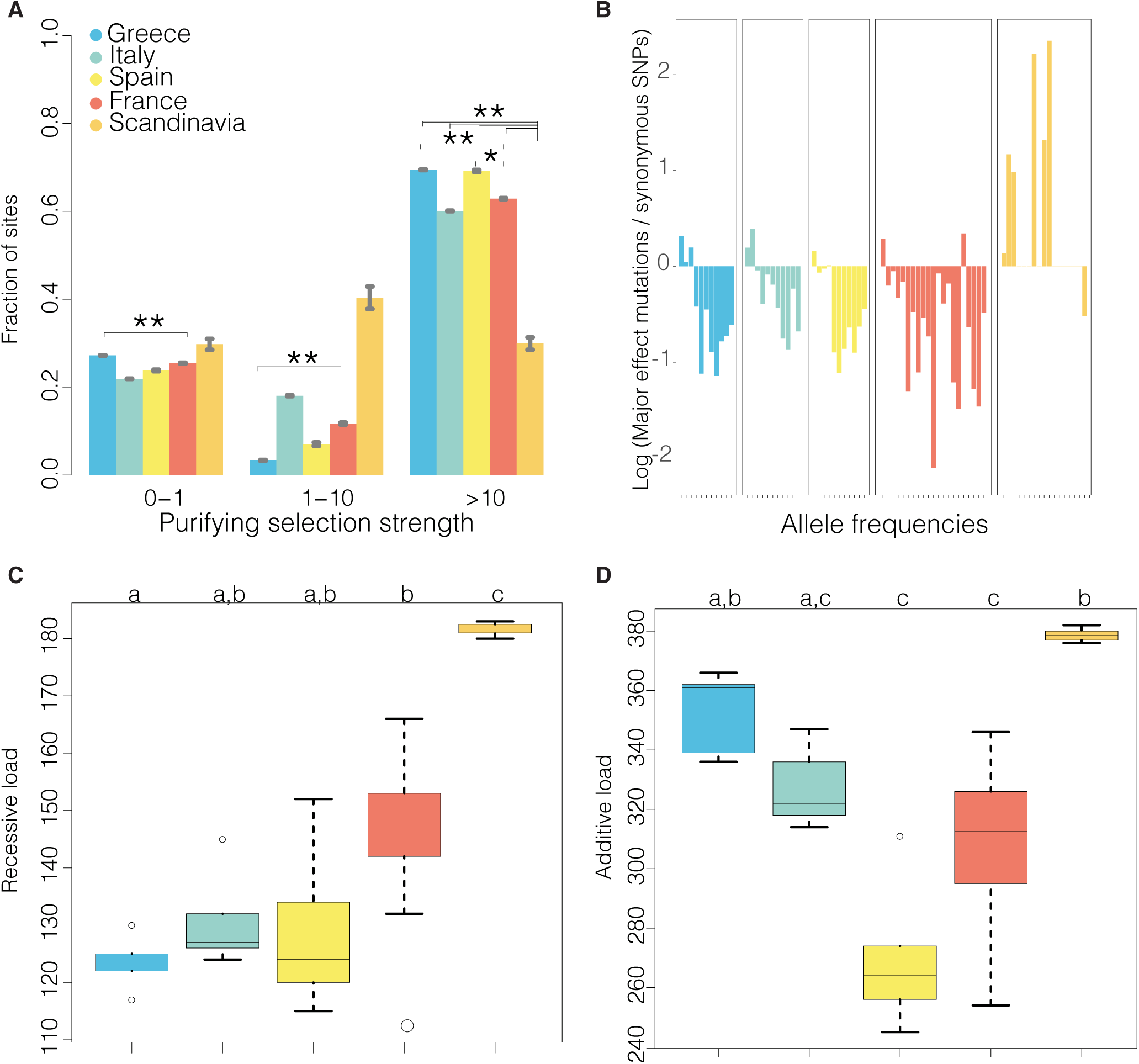
The impact of mating system on purifying selection in *A. alpina*. A. The distribution of fitness effects (DFE) in bins of *N*_*e*_*s* (the product of the effective population size and the selection coefficient) for new nonsynonymous mutations, estimated under model with a stepwise population size change. Error bars show ± 1 standard error. Asterisks indicate significant differences (FDR<0.05) among populations. B. Scandinavian *A. alpina* show an increase in the frequency of derived major effect polymorphism relative to synonymous polymorphism. The figure shows the log ratio of major effect derived allele frequencies to 4-fold synonymous allele frequencies. C. Boxplots of the recessive genetic load for major effect alleles. Letters indicate groups with statistically significant differences (*P*<0.05, Kruskal-Wallis test, post-hoc Dunn test). D. Boxplots of the additive genetic load for major effect alleles. Significance is indicated as in C.

In contrast to results for mixed-mating populations, several lines of evidence suggest that selection against deleterious alleles is compromised in highly self-fertilizing Scandinavian *A. alpina*. First, there was a strong difference in the DFE of nonsynonymous mutations, with a strong, significant reduction in selection against strongly deleterious nonsynonymous mutations (*N*_*e*_*s*>10) in Scandinavia compared to all other regional populations (Fig. 2A). Thus, DFE analyses suggest that the Scandinavian population could be accumulating strongly deleterious nonsynonymous mutations.

To follow up on this observation, we used the closely related species *Arabis montbretiana* as an outgroup to polarize alleles as ancestral or derived. We then examined the allele frequency distribution of derived alleles, and found that derived alleles with a major effect on gene integrity (see Methods) were found at a markedly higher frequency relative to derived 4-fold synonymous alleles in Scandinavia compared to all other regional populations (Fig. 2B). Because selected and neutral allele frequency spectra can be differently affected by demographic changes even in the absence of an actual change in selection (39-41), we considered two additional proxies for genetic load. First, we estimated the average number of homozygous derived major-effect genotypes (Fig. 2C), which is expected to be proportional to genetic load if deleterious mutations act recessively, and then we estimated the average number of derived major-effect alleles (Fig. 2D), which is proportional to genetic load if deleterious mutations have additive effects (42). According to both statistics, the highly selfing Scandinavian regional population had an elevated genetic load compared to mixed-mating populations, and outcrossing and mixed-mating populations were not different from each other (Fig. 2C-D). In addition, the number of fixed derived major effect variants in Scandinavia was highly inflated (∼70% increase) compared to both the outcrossing, and mixed mating populations (SI Appendix, Table S4). A similar pattern was seen for derived nonsynonymous variants and when considering each geographical population separately (SI Appendix, Fig. S3; Fig. S4). Overall, this suggests that the highly selfing *A. alpina* from Scandinavia are accumulating deleterious mutations, whereas mixed-mating and outcrossing populations have similar levels of genetic load.

### A recent bottleneck and selfing explain polymorphism in the Scandinavian population

Theory predicts that high levels of self-fertilization should result in effective purging of recessive deleterious variation, unless selfing is associated with strong reductions in the effective population size (16). Given that we observed no evidence for purging and a strong reduction in genetic variation in Scandinavian *A. alpina*, we asked whether this could be due to a bottleneck, or if increased background selection due to selfing is sufficient to explain the reduction of diversity.

Previous phylogeographic analyses have shown that *A. alpina* likely originated in Asia Minor and subsequently spread westward about 500 kya (28, 30). Population genetic analyses have identified Central European populations as the most likely source for Scandinavian *A. alpina* populations (29), but so far no explicit demographic model has been fit to estimate the timing or demographics of colonization of Scandinavia by *A. alpina*. For this purpose, we used a maximum likelihood-based approach to estimate the parameters of a demographic model of colonization of Scandinavia using a two-dimensional site frequency spectrum (2D-SFS) based on a scattered sample of individuals from Central Europe and the highly selfing Scandinavian *A. alpina* (see Methods). For analyses of demographic history, we used a set of 12,967 SNPs in intergenic regions with low gene density and high recombination rate, which are expected to be less affected by linked selection and thus useful for demographic inference (12, 43). According to our best-fit model, the split between Central Europe and Scandinavia occurred approximately 20 kya and was associated with a prolonged bottleneck (Fig. 3A; SI Appendix, Fig. S5).

**Figure 3.**
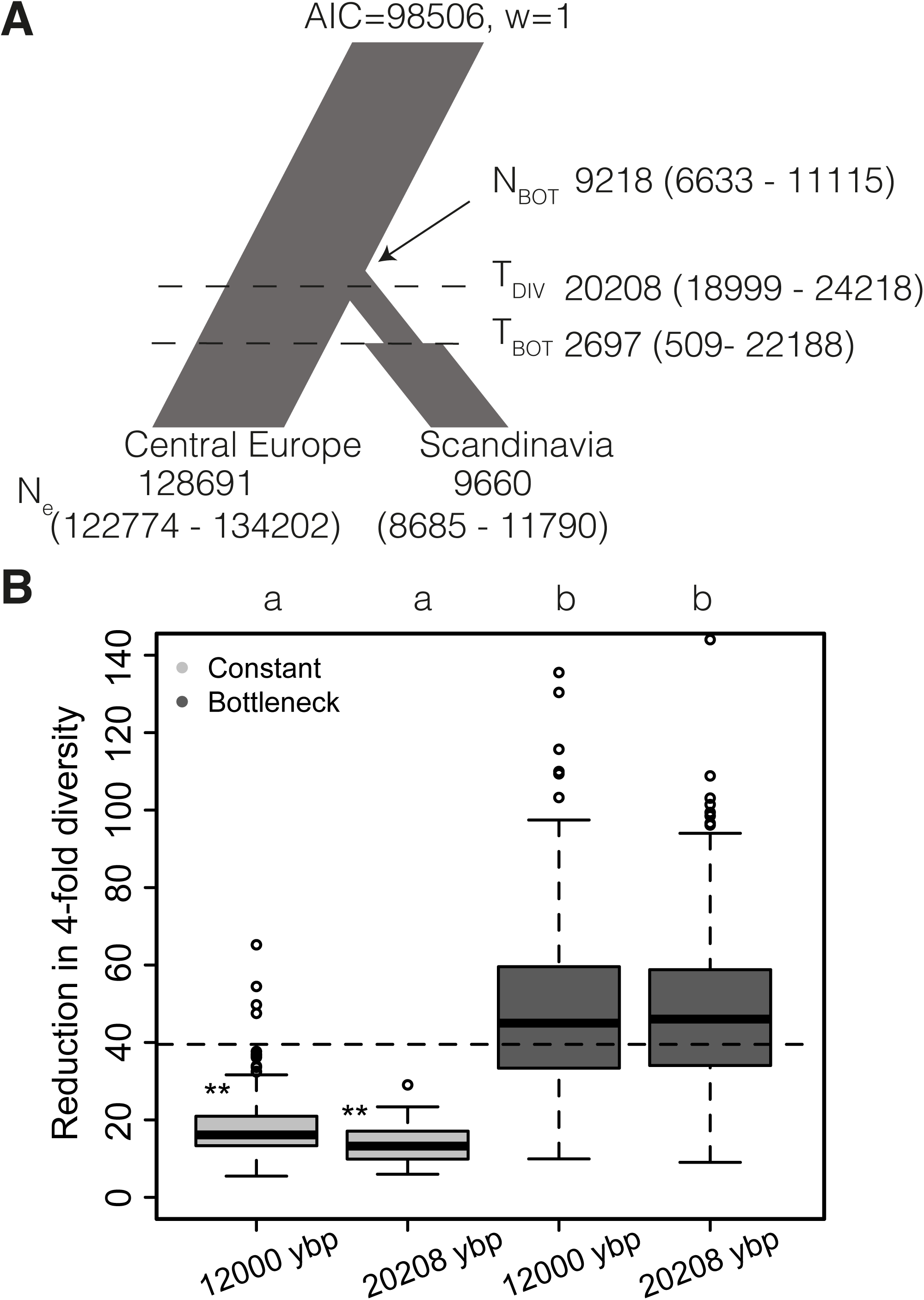
A recent bottleneck and selfing explain the reduction of polymorphism in Scandinavia. A. Schematic showing the best-fit demographic model of the colonization of Scandinavia from an ancestral Central European population. Estimated times are given in years before present (ybp), assuming a generation time of 1.5 years. B. Background selection alone does not explain the reduction in diversity in Scandinavian *A. alpina.* Boxplots show the ratio of synonymous polymorphism between an outcrossing population and a 90% selfing population experiencing either a constant population size or a 10-fold bottleneck, with the two populations diverging either 12,000 ybp or 20,208 ybp. The dashed line indicates the observed ratio of synonymous polymorphism in a scattered sample from Central Europe to that in the Scandinavian regional population. Letters at the top indicate significant difference between models (Mann-Whitney test *P*<0.001). Asterisks represent the significance level of a test of whether the observed neutral diversity reduction is greater than expected based on 300 simulations.

We conducted population genetic simulations to assess whether the reduction in diversity in Scandinavia could be explained by a stronger impact of background selection due to selfing, without additional demographic changes. These simulations, which used realistic settings for *A. alpina* genome structure, including variation in recombination rate, gene density and mutation rates, and a realistic distribution of negative fitness effects (see Methods), show that models that incorporate a transition to selfing but no demographic change cannot explain the reduction in diversity we observe in the Scandinavian population (*P* = 0.013 based on 300 simulations), whereas a model which includes a 10-fold bottleneck and a transition to selfing is consistent with the observed reduction in genetic diversity (*P* = 0.58 based on 300 simulations, Fig. 3B). These conclusions are robust if we assume a more recent split between the Central European and the Scandinavian population, as late as 12,000 years ago (Fig. 3B). These demographic modelling results suggest that a postglacial colonization bottleneck reduced diversity and affected the impact of natural selection in Scandinavian *A. alpina*.

## Discussion

We have used population genomic analyses to investigate effects of mating system and demographic history on genetic variation and purifying selection in the arctic-alpine crucifer species *A. alpina*. Our results show that populations with intermediate levels of self-fertilization maintain genetic variation and experience similar levels of purifying selection as outcrossing populations. We further find that highly selfing Scandinavian *A. alpina* experience relaxed purifying selection, most likely as a result of having undergone a severe bottleneck, the timing of which is consistent with postglacial recolonization of Northern Europe. Our results suggest a strong effect of demography on the impact of purifying selection in selfing populations of this species, and demonstrate that intermediate levels of outcrossing can allow populations to avoid the negative population genetic consequences of self-fertilization.

The fact that mixed-mating populations maintained similar levels of genetic diversity as outcrossing populations suggests that the loss of self-incompatibility in *A. alpina* was not associated with a recent and strong bottleneck. Empirical studies in other plant species have frequently found higher diversity in outcrossing and mixed-mating species relative to highly self-fertilizing species (7, 44), in good agreement with our results. Taken together these findings agree well with the expectation that levels of diversity should be higher in outcrossing than in self-fertilizing populations (6).

We detected no evidence for purging of deleterious alleles in mixed-mating *A. alpina* populations. This suggests that, in contrast to expectations from single-locus theory (16), strongly deleterious alleles may not be rapidly purged as a result of partial selfing in *A. alpina*, perhaps due to selective interference (5). While our power to detect purging may be limited if strongly deleterious recessive mutations are rare, a recent simulation-based study showed that the analysis method used here can detect the signal of purging following shifts to higher selfing rates (38). Indeed, no strong differences between outcrossing and mixed-mating populations were found for the distribution of fitness effects of new nonsynonymous mutations, derived allele frequencies of major-effect alleles or for two different proxies of genetic load that assume different dominance levels of deleterious alleles. Our results thus suggest that the genome-wide impact of purifying selection is similar in outcrossing and mixed-mating populations.

While there is a dearth of genome-wide studies contrasting purifying selection in outcrossing and mixed-mating plants, one previous study also found modest differences in purifying selection between outcrossing and mixed-mating species (24). These observations agree with simulation-based results that suggest a low degree of outcrossing is sufficient to maintain efficient purifying selection (23). Thus, mixed-mating populations may avoid some of the negative evolutionary genetic effects associated with high selfing rates, which could imply that population genetic effects contribute to the stable maintenance of intermediate outcrossing rates (5). However, ultimately, direct estimates of inbreeding depression are needed to fully elucidate the maintenance of mating system variation in *A. alpina*.

In contrast to the lack of differences in purifying selection between outcrossing and mixed-mating populations, we found strong evidence for accumulation of deleterious mutations in highly self-fertilizing Scandinavian *A. alpina*. Indeed, our DFE analyses indicate less efficient selection against strongly deleterious alleles, and we observe an increase in the relative allele frequency and fixed derived major-effect mutations, as well as elevated additive and expressed genetic load in Scandinavian *A. alpina*. These results suggest that deleterious alleles are not efficiently selected against and have been able to increase in frequency in this population. In the context of human population history, theory and simulations have shown that range expansions (45) or strong and extended bottlenecks (39, 41, 49) can lead to increased genetic load. It has previously been shown that genetic variation is strongly reduced over a vast geographical area representing the northern part of the distribution of *A. alpina* (29). Our demographic inference and population genetic simulations suggest that this reduction of genetic variation is most likely a result of postglacial colonization bottlenecks. Thus, bottlenecks associated with range expansion appear to have strongly reduced the effective population size and increased the impact of drift in the Northern part of the species range, causing accumulation of deleterious variants in Scandinavian *A. alpina*. The impact of demographic history on genetic load is currently a strongly debated topic in human population genetics, where studies on the effect of the out-of-Africa bottleneck on genetic load have come to different conclusions depending on the statistics they applied (39, 46-48). Here, we extend these studies to a plant species and document increased accumulation of deleterious mutations in bottlenecked Scandinavian *A. alpina*.

Increased background selection can lead to sharply reduced genetic diversity in highly selfing populations (7, 12), but our forward population genetic simulations show that this effect alone cannot explain the reduction in diversity in Scandinavia, whereas a model with both selfing and a population size reduction is consistent with the observed reduction in diversity. At present, however, we cannot rule out a contribution of positive selection during range expansion to reduced diversity in the Scandinavian *A. alpina* population. Indeed, reciprocal transplant experiments have documented adaptive differentiation between Spanish and Scandinavian populations of *A. alpina* (49). While elevated load can be associated with enhanced local adaptation following a range expansion (50), this raises the possibility that some of the increase in frequencies of major-effect variants in Scandinavia may have been directly or indirectly driven by positive selection. However, we believe that it is unlikely to result in the genome-wide signature of relaxed purifying selection that we observe in Scandinavian *A. alpina*. Empirical identification of genomic regions responsible for local adaptation and population genomic studies using larger sample sizes will be needed to explore the genomic impact of positive selection.

Here, we have investigated the impact of demographic history and mating system on genomic patterns of variation in *A. alpina*. Our results show that mixed-mating populations maintain genetic variation and purifying selection at similar levels as outcrossing populations. In contrast, we find an increase in genetic load in a highly self-fertilizing population, most likely as a result of demographic effects associated with postglacial range expansion. Our results are important for a more general understanding of the impact of mating system and demographic history on genomic variation and selection in plants.

## Methods

### Data and sequencing

We performed paired-end (100 bp) whole-genome resequencing of 38 *Arabis alpina* individuals from Europe (see SI Appendix, Table S1), utilizing libraries with an insert size of 300-400 bp (see SI Appendix, SI Text) on an Illumina HiSeq 2000 instrument (Illumina, San Diego, CA, USA). We obtained a total of 10,079 Gbp and ∼245 Gbp (QC > 30) per sample on average with a mean coverage of 26X ranging from 16X to 45X.

### Quality assessment, trimming, genotype calling and filtering

Adapter trimmed reads were mapped to the *A. alpina* V4 reference genome assembly (30) using BWA-MEM v0.7.8 (51), and duplicate free BAM alignment files were further processed using the Genome Analysis Toolkit (GATK v.3.4.0; 52) (see SI Appendix, SI Text). The *A. alpina* genome assembly is exceptionally enriched for repetitive elements relative to previously sequenced Brassicaceae relatives (31) and so we employed a variety of hard and custom filtering techniques (see SI Appendix, SI Text) to avoid calling SNPs in regions of the genome that putatively represent copy number variants. After applying all filters the dataset was composed of 1,514,615 SNPs and 43,209,020 invariant sites amenable to further analysis.

### Inference of population structure and population genetic analyses of selection

Population structure was inferred using a combination of principal component analysis (PCA), and Bayesian clustering analysis using both fastSTRUCTURE v1.0 (34) and TESS v3 (35) (SI Appendix, SI Text).

For each of the five regional populations identified by population structure analysis, we obtained estimates of summary statistics (S, π, Tajima’s D) (Table 1, see SI Appendix, SI Text). We estimated the distribution of fitness effects (DFE) using DFE-alpha v. 2.15 (37) on folded 4-fold and 0-fold site frequency spectra (SFS) under a stepwise population size change model (SI Appendix, SI Text). DFE estimates were compared among regional populations based on 200 bootstrap replicates.

### Major effect mutations and genetic load

Presence and frequency of major effect mutations, i.e. loss of start and stop codons; gain of stop codons and changes in splice sites were calculated per population using snpEFF v4.2 (53). To avoid reference-biased inference of alternate alleles we polarized the SNPs using the *A. montbretiana* genome assembly (ASM148412v1, 54) as an outgroup (SI Appendix, SI text). As a proxy for genetic load, we estimated the average number of derived nonsynonymous or major-effect homozygous genotypes (41, 46), and the average number of derived alleles for nonsynonymous and major-effect alleles (42) for each regional population. In addition, we counted the total number of fixed derived nonsynonymous or major-effect alleles for each regional population.

### Demographic modelling and simulations

We conducted demographic inference in the software fastsimcoal2 v. 2.5.2.21 (55). To estimate parameters associated with the origin of Scandinavian *A. alpina*, we tested two demographic models (SI Appendix, Fig. S5), using two-dimensional joint SFS (2D-SFS) based on a scattered sample from central Europe and the Scandinavian population (SI Appendix, SI Text). We used 12,967 intergenic sites, a mutation rate of 7 × 10^-9^ and a generation time of 1.5 years. We used forward simulation in SLiM2 v2.1 (56) to assess the impact of demography and selection associated with a shift to selfing on genetic diversity in the Scandinavian population under four demographic models with varying bottleneck severity and population split time (SI Appendix, SI Text).

## Acknowledgements

TS thanks Cindy Canton for help with plant care and extractions. Computations were performed on resources provided by SNIC through UPPMAX under Projects snic2014-1-194, b2013022 and b2013237. This work was funded by grants from the Swedish Research Council to JÅ and TS, by a grant from the DFG through SPP1529 to GC, and from SciLifeLab to TS.

